# Rolling Circle Amplification is a high fidelity and efficient alternative to plasmid preparation for the rescue of infectious clones

**DOI:** 10.1101/2020.06.22.165241

**Authors:** Jeffrey M. Marano, Christina Chuong, James Weger-Lucarelli

**Affiliations:** Department of Biomedical Sciences and Pathobiology, Virginia Tech, VA-MD Regional College of Veterinary Medicine, Blacksburg, VA, USA; Translational Biology, Medicine, and Health Graduate Program, Virginia Tech, Blacksburg, VA, United States

## Abstract

Alphaviruses (genus *Alphavirus*; family *Togaviridae*) are a medically relevant family of viruses that include chikungunya virus, Eastern equine encephalitis virus, and the emerging Mayaro virus. Infectious cDNA clones of these viruses are necessary molecular tools to understand viral biology and to create effective vaccines. The traditional approach to rescuing virus from an infectious cDNA clone requires propagating large amounts of plasmids in bacteria, which can result in unwanted mutations in the viral genome due to bacterial toxicity or recombination and requires specialized equipment and knowledge to propagate the bacteria. Here, we present an alternative to the bacterial-based plasmid platform that uses rolling circle amplification (RCA), an *in vitro* technology that amplifies plasmid DNA using only basic equipment. We demonstrate that the use of RCA to amplify plasmid DNA is comparable to the use of a midiprepped plasmid in terms of viral yield, albeit with a slight delay in virus recovery kinetics. RCA, however, has lower cost and time requirements and amplifies DNA with high fidelity and with no chance of unwanted mutations due to toxicity. We show that sequential RCA reactions do not introduce mutations into the viral genome and, thus, can replace the need for glycerol stocks or bacteria entirely. These results indicate that RCA is a viable alternative to traditional plasmid-based approaches to viral rescue.

**Importance:** The development of infectious cDNA clones is critical to studying viral pathogenesis and for developing vaccines. The current method for propagating clones in bacteria is limited by the toxicity of the viral genome within the bacterial host, resulting in deleterious mutations in the viral genome, which can only be detected through whole-genome sequencing. These mutations can attenuate the virus, leading to lost time and resources and potentially confounding results. We have developed an alternative method of preparing large quantities of DNA that can be directly transfected to recover infectious virus without the need for bacteria by amplifying the infectious cDNA clone plasmid using rolling circle amplification (RCA). Our results indicate that viral rescue from an RCA product produces a viral yield equal to bacterial-derived plasmid DNA, albeit with a slight delay in replication kinetics. The RCA platform, however, is significantly more cost and time-efficient compared to traditional approaches. When the simplicity and costs of RCA are combined, we propose that a shift to an RCA platform will benefit the field of molecular virology and could have significant advantages for recombinant vaccine production.

## 1. Introduction

RNA viruses produce significant disease in humans and animals, highlighted by the current outbreak of severe acute respiratory syndrome coronavirus 2 (SARS-CoV-2) [1]. Infectious cDNA clones of these viruses are necessary molecular tools to understand viral biology since they facilitate the study of single-nucleotide polymorphisms [2] and enable the insertion of reporter proteins to study virus replication or cell tropism [3]. cDNA clones have also been instrumental in developing vaccines for RNA viruses, notably CYD-TDV (Dengvaxia) [4] and TAK-003 (Takeda)[5], both of which are tetravalent chimeric vaccines against dengue virus.

Alphaviruses (genus *Alphavirus*; family *Togaviridae*) are a group of small, enveloped, medically relevant positive-sense RNA viruses with genomes of 11-12 kilobases in length [6]. Examples of medically relevant alphaviruses include several arthropod-borne viruses, or arboviruses, such as chikungunya virus, Ross River virus, Eastern equine encephalitis, and the emerging Mayaro virus [7]. Additionally, alphavirus genomes are relatively easy to manipulate and have been used as expression vectors for foreign proteins [8, 9].

Typically, the propagation of infectious cDNA clones before viral rescue requires the generation of high concentration plasmid stocks from bacteria, which is not only cumbersome and time-consuming but also presents an opportunity for the introduction of unwanted mutations during amplification in bacteria. Bacterial instability of viral genomes has been reported for flaviviruses [10], alphaviruses [11], and coronaviruses [12]. The cause of this is likely cryptic prokaryotic promoters, which results in the expression of viral proteins inside of the bacteria, which due to their toxicity, can lead to the selection of plasmids with deletions, mutations, or recombination with reduced bacterial toxicity [13-15]. There are several critical points within the plasmid-based rescue workflow where deletion or mutations can occur (Figure 1). These include the transformation of the plasmid into the bacteria, the selection of colonies from the agar plate, and the propagation of the colony in liquid culture. While deletions are easy to identify in plasmids by restriction enzyme digestion, mutations can only be determined by whole-genome sequencing, which is costly and laborious. Furthermore, even synonymous changes can have profound impacts on viral replication [16-18] and should be avoided in cDNA clones. These unwanted changes to the viral genome can confound experimental results and, therefore, necessitate sequencing of the full viral genome every time new plasmid stocks are generated, a time-consuming and expensive task. Thus, removing the need for the bacterial host to maintain and propagate infectious cDNA clone plasmids would simplify the process and remove the possibility of deleterious bacterial-derived mutations.

**Fig. 1:**
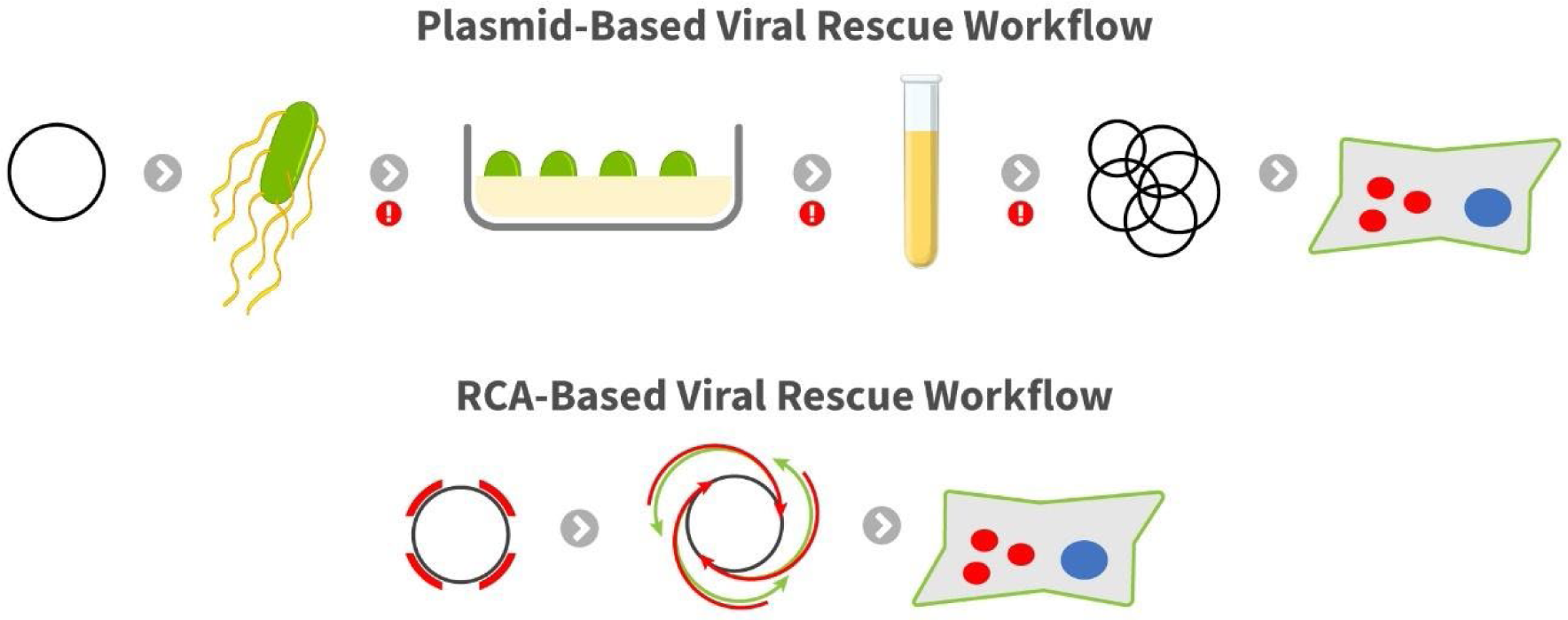
Comparison of plasmid- and RCA-based workflows for viral rescue. The plasmid-based system involves the transformation of a plasmid into bacteria. The bacteria are then selected and propagated using antibiotic enriched media, and the plasmid is purified from the bacteria and transfected into the cell type of interest. The red exclamation points indicate points during the workflow where mutations and error can be introduced or enhanced. The RCA-based system involves amplification of the plasmid using random hexamers to produce the hyperbranched product. This product can then be directly transfected into cells.

In this report, we describe an alternative to bacterial-based growth of infectious cDNA clones—*in vitro* amplification using rolling circle amplification (RCA). RCA is an isothermal, high yield method of DNA amplification [19] that uses a highly processive polymerase that can amplify DNA over 70kb [20]. Importantly, the enzyme replicates DNA with high fidelity due to its 3’—5’exonuclease—or proofreading—activity [21]. We show that peak virus yields were similar following RCA- and plasmid-derived virus rescue in several cell lines and that small amounts of the RCA product could be used to rescue virus successfully. Finally, we showed that we could further amplify an RCA product through additional rounds of RCA without the introduction of unwanted mutations, thereby allowing a simple, cheap, and high-fidelity means to propagate infectious cDNA clone plasmids. RCA-launched infectious cDNA clones represent a technical improvement in rescuing viruses without the need for bacteria.

## 2. Materials and Methods

### Cell Culture

Vero (*Cercopithecus aethiops* kidney epithelial cells, ATCC® CCL-81™), BHK-21 clone 13 (baby hamster kidney fibroblasts, ATCC® CCL-10™), and HEK293T (human embryonic kidney cells, ATCC® CRL-11268™) cell lines were maintained at 37°C in 5% CO2 using Dulbecco’s modified Eagle’s medium (DMEM) supplemented with 5% fetal bovine serum (FBS), 1% nonessential amino acids, and 0.1% gentamicin. Plaque assays were performed as previously described [17] except that plates were either fixed for two days (1% gum tragacanth overlay-MP BIOMEDICALS catalog 0210479280) or three days (1.5% methylcellulose overlay-Spectrum Chemical catalog ME136-100GM) post-infection.

### Rescue of infectious cDNA clones

We performed transfections in 24 well plates at 60-80% confluency using the JetOptimus (Polyplus) DNA transfection reagent per manufacturer’s instructions. Briefly, we mixed 200 µl of JetOptimus buffer with the DNA concentration of interest. JetOptimus reagent was then added at a ratio of 1 µl per 1 µg of DNA and incubated at room temperature for 10 minutes. The transfection mix was then added dropwise into the wells containing cells. We collected the supernatant at different time points based on the experiment: each day for three days for the growth curves in multiple cell lines, each day for two days for the RCA input comparison and the kit comparison, and two days post-transfection for the sequential RCA experiment. Viral titer was then determined using plaque assays on Vero cells.

### Plasmid Preparation

We used an infectious cDNA clone of Mayaro virus strain TRVL 4675, which has previously been described [22], for our plasmid control. The plasmid was initially transformed into NEBstable electrocompetent cells. Cells were incubated for 16 hours at 30°C and then 24 hours at room temperature. Colonies were picked and incubated in Lennox Broth (LB) supplemented with 25 µg/ml of carbenicillin for 16 hours. We extracted DNA using both Promega PureYield miniprep kit, for verification, and Zymo Midiprep Kit, for transfection. We verified the plasmids using endonuclease digestion and gel electrophoresis and transfected the samples to ensure the infectivity of the clone. DNA concentration was determined using Invitrogen’s Qubit 1x dsDNA HS kit.

### RCA Protocols

For the SuperPhi RCA Premix Kit with Random Primers (Evomics catalog number PM100), 1 µl of 1 ng/µl of plasmid DNA was mixed with 4 µl of sample buffer while the thermocycler was preheated to 95°C. The mixture was incubated at 95°C for 1 minute and then rapidly cooled to 4°C. 5 µl of 2x SuperPhi Master Mix was then mixed with the sample, which was then incubated for 16 hours at 30°C before polymerase inactivation at 65°C for 10 minutes. For the GenomiPhi V3 DNA Amplification Kit (GE Healthcare), 1 µl of 10 ng/µl of plasmid DNA was mixed with 9 µl of molecular grade water and 10 µl of denaturation buffer. At the same time, the thermocycler was preheated to 95°C. The mixture was incubated for one minute at 95°C and then rapidly cooled to 4°C. 20 µl of the denatured template was then added to the lyophilized reaction cake containing enzymes, dNTPs, and buffers and thoroughly mixed by pipetting. Samples were incubated at 30°C for 90 minutes, and then the enzyme was inactivated at 65°C for 10 minutes. RCA product concentration was determined using Qubit after a 200-fold dilution in molecular grade water. Amplification of the plasmid was confirmed using endonuclease digestion and gel electrophoresis. Dilutions were performed using molecular grade water. To generate the RCA passages, an initial RCA was performed using the SuperPhi protocol as described above and validated using gel electrophoresis. 1 µl of RCA product was then used as the template for a subsequent 10 µl RCA reaction. We then repeated the process of RCA for a total of three passages.

### Sequencing Protocol

RCA products were Sanger sequenced at the Genomics Sequencing Center at Virginia Tech. RCA products were diluted to 100 ng/µl to prepare them for sequencing. 1 µl of diluted RCA product was mixed with 3 µl of 1 µM primer and 9 µl of molecular grade water. Resulting reads were aligned using SnapGene® 5.0.7 software (GSL Biotech).

### Statistical Analysis

Statistics were performed using GraphPad Prism 8 (San Diego, CA). Two-way ANOVA tests were performed using Sidak’s corrections for multiple comparisons for the comparison of titers in different cell lines. For the comparison of RCA kits, two-way ANOVA tests were performed using Dunnett’s correction for multiple comparisons against the plasmid control. A one-way ANOVA was performed using Sidak’s correction for multiple comparisons against the plasmid control for the sequential RCA test.

## 3. Results and Discussion

### Peak virus yields are similar for RCA- and plasmid-derived virus in different cell lines

To examine the efficacy of RCA for viral rescue, we chose three cell lines for transfection: Vero, HEK293T, and BHK21 Clone 13 cells (BHK21). We selected these cell lines due to their widespread use in virus rescue [23, 24]. We used a CMV-driven Mayaro virus infectious cDNA clone as both the template for our RCA reactions and as our plasmid control for the transfections (Fig. 2). Following transfection, we collected virus each day until 90% of the cells showed cytopathic effect (CPE), which occurred by day three in all cases. On the first day post-transfection, viral titer in the plasmid transfection was significantly higher compared to the RCA product in both BHK21 and HEK293T cells (p=0.0002 and p= 0.0007, respectively). No difference was observed in Vero cells (p=0.0961). There was no significant difference in any cell line (Vero p= 0.9445, BHK21 p=0.2937, HEK293T p=0.0599) two days post-transfection, the peak of virus replication for all cell lines. At three days post-transfection, there was no difference between viral titers produced by RCA and plasmid in Vero and BHK21 cells (p=0.1736 and p=0.6140, respectively). However, the titer of RCA product transfection was significantly higher than the plasmid titer in HEK293T cells (p=0.0027).

**Fig. 2:**
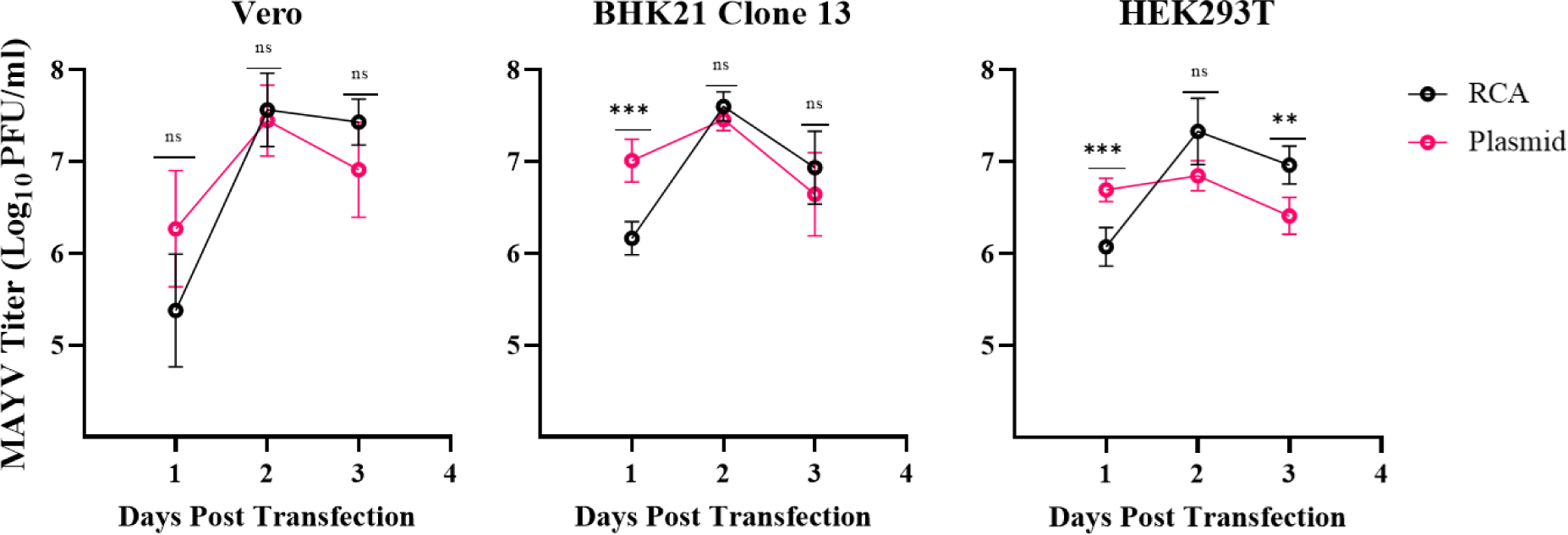
Comparison of viral titers produced by either plasmid or RCA in various cell lines. The kinetics of plasmid and RCA techniques to produce virus were assessed in Vero, BHK21 Clone 13, and HEK293T cells. Cells were transfected in triplicate with either 500 ng of Evomics SuperPhi RCA or plasmid DNA in triplicate. The experiment was done in two independent biological replicates. The supernatant was collected each day post-transfection until cells reached 90% CPE for plaque assay. Error bars represent the standard deviation from the mean. Statistical analysis was performed using two-way ANOVA with *ad hoc* Sidak’s correction for multiple comparisons (ns P > 0.05, ** P ≤ 0.01, *** P ≤ 0.001).

The above results demonstrate two critical features of using RCA for viral rescue: similar replication kinetics—albeit with a slight delay—and identical peak yield. In all the cell lines, the peak viral titer occurred on the second-day post-transfection for both plasmid and RCA product transfection. These data demonstrate that RCA is equivalent to plasmids in their ability to rescue virus using standard transfection conditions in terms of viral yield. A hypothesis for the delay in viral production seen in BHK and HEK293T cells is that the complex structure of RCA products (i.e., branched RCA molecules) compared to plasmid DNA may produce steric inhibition and delay transcription from the CMV promoter by RNA polymerase II [19].

### Peak viral titer is not dependent on DNA input or RCA kit

Since the kinetics of virus recovery were similar for all cell lines, and since peak titers were observed two days post-transfection, we only used Vero cells and only sampled on the first- and second days following transfection for all future studies. We next sought to determine whether transfections with different RCA product input concentrations would result in efficient virus rescue. To that end, we transfected Vero cells with a range of RCA inputs produced using the Evomics SuperPhi kit (Fig. 3). One-day post-transfection, 100 ng of RCA resulted in a decreased viral titer compared to a plasmid input of 500 ng (p = 0.0002). We observed no differences in any of the other input concentrations. As in the above experiment, by two days post-transfection, RCA and plasmid titers were the same (100 ng SuperPhi p= .9275, 250 ng SuperPhi p=.9991, 500 ng p>.9999, 1000 ng p=.9952). These results indicate that the transfection of RCA product is robust and can tolerate a wide variety of input concentrations without altering peak viral yield.

**Fig. 3:**
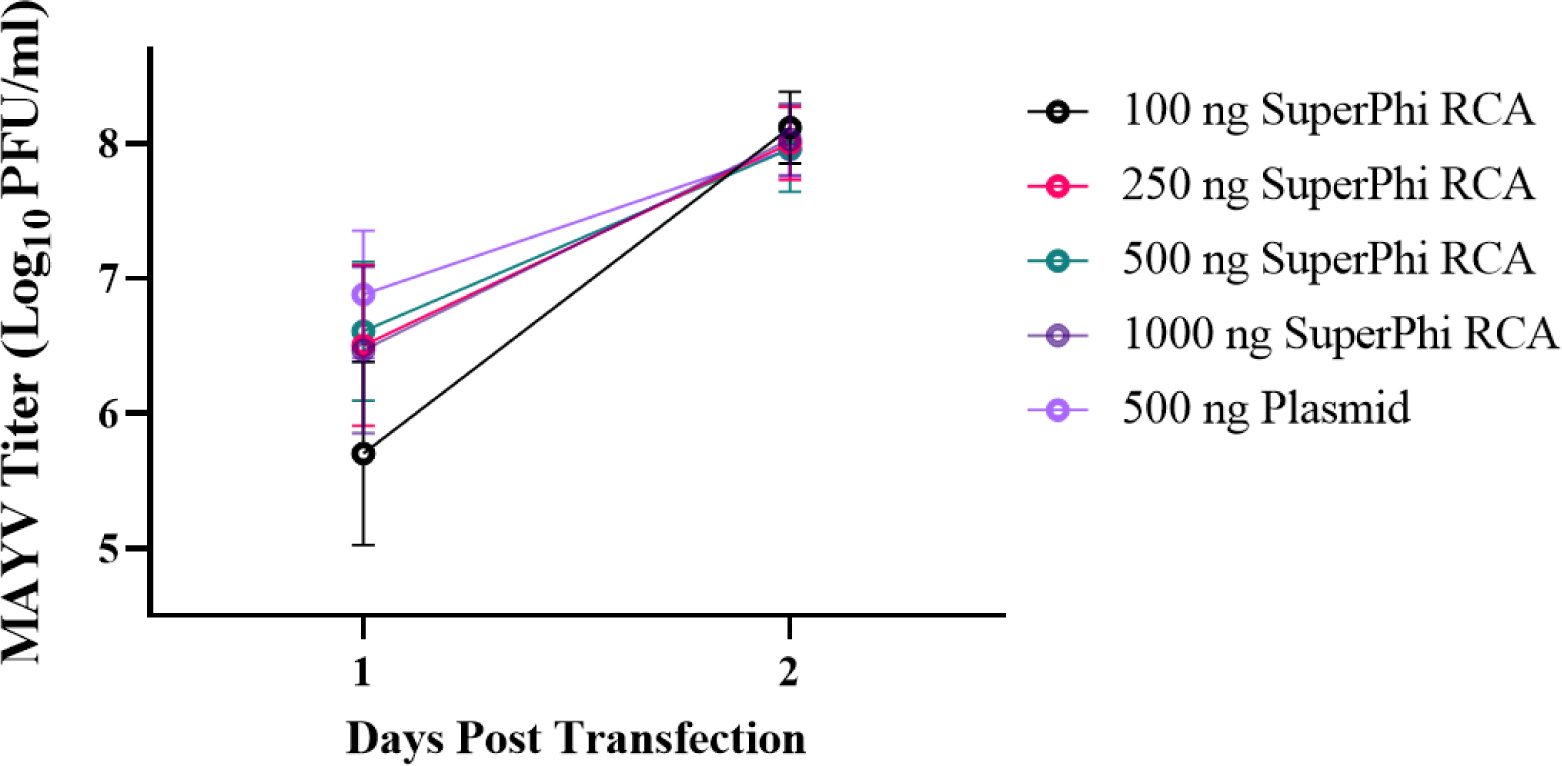
Assessing the effects of RCA input on the resulting viral titer. The effect of RCA input on viral production kinetics was examined using Vero cells. Cells were transfected in triplicate with 100 ng, 250 ng, 500 ng, or 1000 ng of Evomics SuperPhi RCA or 500 ng of the plasmid. The supernatant was collected at one and 2-days post-transfection for plaque assay. Error bars represent the standard deviation from the mean. Statistical analysis was performed using two-way ANOVA with *ad hoc* Dunnett’s correction for multiple comparisons.

To ensure that the above results were not restricted to a specific RCA kit, Vero cells were transfected in triplicate in two independent replicates with RCA product produced using both the Evomics SuperPhi Kit and the GE GenomiPhi Kit or plasmid DNA (Fig. 4). One day post-transfection, the viral titers produced from both 250 ng and 500 ng of GenomiPhi RCA products were lower than the titers produced by plasmid (p = 0.0008 and p = 0.0005, respectively). There was no significant difference between the SuperPhi samples and the plasmid samples one-day post-transfection (250 ng p=.4915, 500 ng p=.4490). All titers were the same two days post-transfection compared to plasmid rescue (250ng SuperPhi p=.9997, 500 ng SuperPhi p=.9748, 250 ng GenomiPhi p=.9966, 500 ng GenomiPhi p=.9927.) Thus, peak viral yields or recovery kinetics of RCA products are not dependent on the RCA kit.

**Fig. 4:**
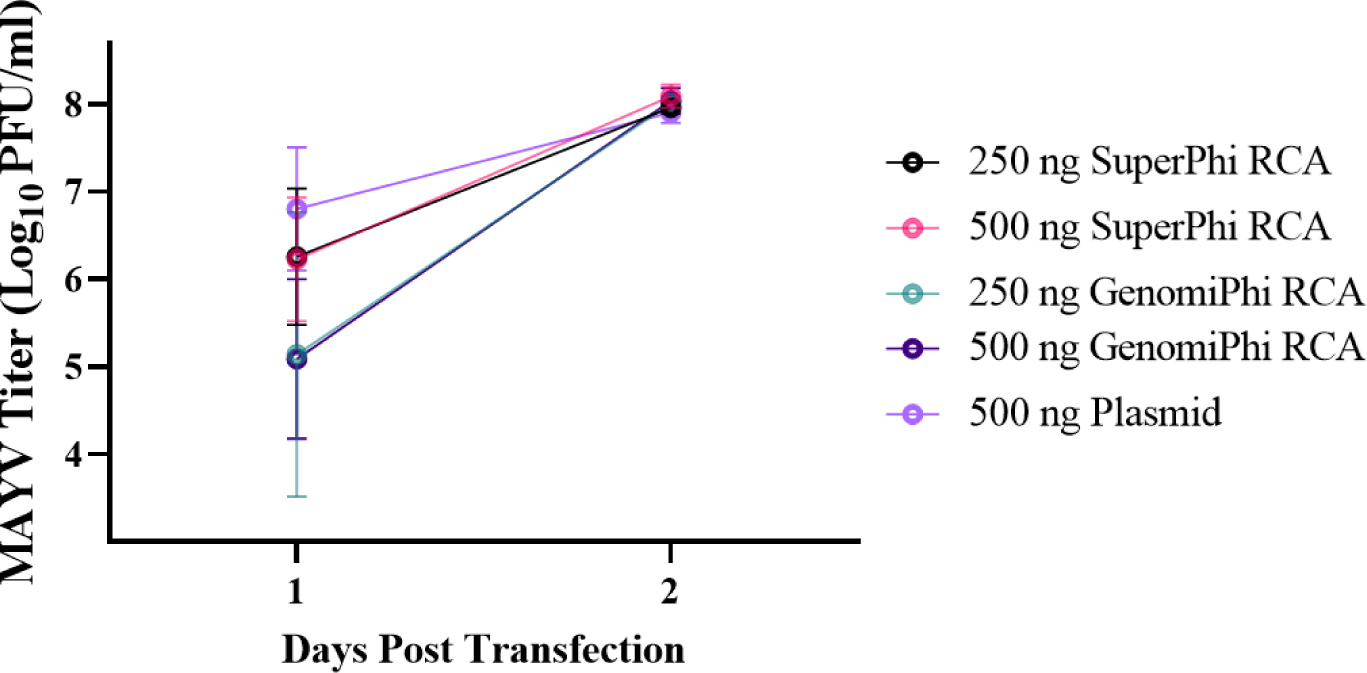
Assessing the effects of RCA kits on resulting viral titer. The effect of RCA kits on viral production kinetics was examined using Vero cells. Cells were transfected with 250 ng or 500 ng of either Evomics SuperPhi or GE GenomiPhi RCA or 500 ng of plasmid DNA. The supernatant was collected at one and 2-days post-transfection for plaque assay. Error bars represent the standard deviation from the mean. Statistical analysis was performed using two-way ANOVA with *ad hoc* Dunnett’s correction for multiple comparisons against a plasmid control.

### Sequential RCA allows for simple propagation of an infectious cDNA clone without introducing errors in the viral genome

To determine if RCA can further amplify an RCA product without introducing unwanted mutations, an initial RCA was performed using plasmid DNA as a template (subsequently referred to as Passage 0) and amplified three more times. Following the transfection of the different “passages,” we found no significant differences between the viral titer produced in passage 0 RCA DNA and plasmid DNA (p= 0.2518) (Fig. 5). However, we did note a difference between the titers of later RCA passages and plasmid DNA (p = 0.0008, p = 0.0007, and p = 0.0016, respectively). To determine if these differences were due to either mutations or artifacts of repeated amplification, RCA DNA from passage 0 and 3 was sequenced using Sanger Sequencing. The sequences for passage 0, passage 3, and the original plasmid were identical, indicating that no mutations were introduced in the viral genome during repeated RCAs. The likely cause of the reduction in peak viral yield is that the repeated RCAs amplified both specific and non-specific DNA. Amplification of non-specific DNA, which is caused by the concatemerization of the random hexamer primers [25], alters the ratio of specific to non-specific DNA, resulting in a reduction in the amount of target DNA that is transfected. To mitigate the effect of moderate titer reduction with sequential RCA reaction, harvesting virus at a slightly later time or increasing DNA input may be effective. However, these results indicate that RCA products can effectively act as a template for subsequent RCA reactions without introducing unwanted mutations. RCA products can, therefore, substitute the standard glycerol bacterial stock protocol, or repeated bacterial transformations to generate midi- or maxi-prepped DNA. In both bacterial methods, mutations can be introduced during growth and, thus, require sequencing.

**Fig. 5:**
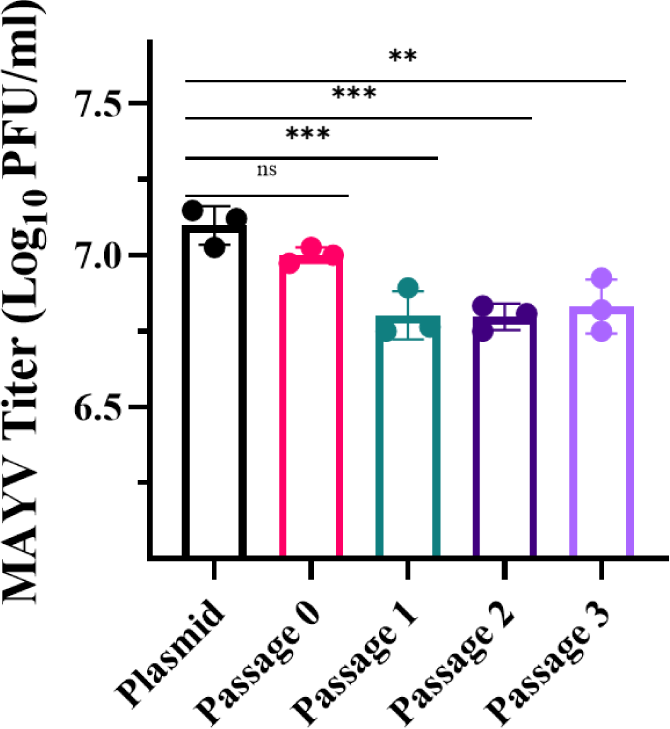
Assessing the effects of repetitive RCA on resulting viral titer. We examined the impact of sequential RCA on the ability to rescue virus in Vero cells. We used the Evomics SuperPhi RCA kit and a sample of midiprepped DNA as a template to generate RCA products (Passage 0). We then used the RCA product as the template for subsequent RCA reactions (Passage 1-3). We transfected the cells using 500 ng of RCA product or 500 ng of plasmid DNA in triplicate. The supernatant was collected 2-days post-transfection for plaque assay. Error bars represent the standard deviation from the mean. Statistical analysis was performed using one-way ANOVA with *ad hoc* Dunnett’s correction for multiple comparisons against the plasmid control (ns P > 0.05, ** P ≤ 0.01, *** P ≤ 0.001).

## 4. Conclusion

Here, we report a simple method to recover infectious virus from a cDNA clone using RCA to amplify a plasmid. We observed that both RCA and plasmid-based transfection produced similar peak viral titers following transfection for several cell lines, using several RCA kits, and when transfecting variable input DNA amounts. Importantly, RCA products can be reamplified by RCA to maintain a DNA record without generating mutations in the viral genome.

The evidence above demonstrated that the RCA platform is equivalent to the plasmid-based platform in terms of viral yield. However, when considering the time and cost to perform these two processes, it is apparent that RCA offers many advantages (Table 1). Using the SuperPhi kit as an example, one RCA reaction costs $3.79 and produces 10 µg of DNA in 16 hours. When using the plasmid approach, first, you would need to screen colonies using endonuclease digestion. Assuming you screen ten colonies using the Promega PureYield™ Plasmid Miniprep System, it will cost $15.70. From there, positive colonies would then be used to inoculate cultures for midiprep. Assuming you midiprep between one and four colonies using the ZymoPURE™ II Plasmid Midiprep Kit, this purification would cost between $9.16 and $36.64. The extracted midiprep DNA would then be used for Sanger sequencing, costing roughly $150 per genome. Therefore, the final cost of the plasmid workflow ranges from $174.86 to $652.34, with a final DNA yield of 80 µg. When comparing cost per µg of DNA, it is apparent that the RCA system is superior to the plasmid approach. RCA reactions cost $0.38/µg, while the plasmid approach costs between $2.19-$8.15/µg. This difference is further emphasized when considering the time to complete the two reactions. An RCA reaction takes 16 hours or less and can be transfected directly while the bacterial approach takes several days and requires sequencing. Given this cost and time analysis, the RCA platform is a more time and cost-efficient method for rescuing viruses and produces similar results, indicating that a shift to the RCA-based approach would simplify viral rescue while saving time and money.

**Table 1:** Comparison of the cost and time requirements for the plasmid and RCA systems. The cost calculations in the table are based on the following kits: Promega PureYield™ Plasmid Miniprep System (catalog A1222), ZymoPURE™ II Plasmid Midiprep Kit (catalog D4201), and Evomics SuperPhi RCA Premix Kit with Random Primers (catalog number PM100). In constructing the cost estimate, it was assumed that ten colonies were selected for screening, and one to four of those colonies were then midiprepped and sequenced.

| Stepwise Cost and Time Analysis of RCA and Plasmid Systems |  |  |  |  |  |
| --- | --- | --- | --- | --- | --- |
| Plasmid |  |  | RCA System |  |  |
| Protocol | Cost | Time | Protocol | Cost | Time |
| Transformation and Miniprep | \$15.70 | 3 days | RCA Amplification | \$3.79 | 16 hours |
| Midiprep | \$9.16 - \$36.64 | 1 day | DNA Yield | 10 µg | |
| Sequencing | \$150 - \$600 | 2 days | Cost per µg | \$.38/µg | |
| Total | \$174.86 - \$652.34 | 6 days | | | |
| DNA Yield | 80 µg |  |  |  |  |
| Cost per µg | \$2.19 - \$8.15/µg | | | | |

The use of RCA-launched expression from RNA polymerase II promoter containing constructs has several potential commercial applications as well, including DNA and recombinant live-attenuated vaccines. Bacterial-derived plasmids have several safety concerns, including endotoxins, transposition of pathogenic elements, and the introduction of antibiotic resistance into the environment [26]. RCA mitigates many of these concerns since the antibiotic resistance marker is not required, and RCA products are free from endotoxin.

This study has two limitations: first, we only used a single MAYV infectious cDNA clone to characterize the RCA rescue system. However, we successfully used RCA to amplify and rescue a variety of infectious cDNA clones, including Zika virus, chikungunya virus, Usutu virus, and Sindbis virus. We anticipate that this system can be used for other positive-sense RNA virus cDNA clones and likely negative-strand viruses as well. Second, we only tested viral rescue from a clone driven by a cytomegalovirus (CMV) promoter. Using a CMV promoter allows for simple transfection of small amounts of plasmid DNA without the need for extra reagents to produce viral RNA or potential issues with unwanted mutations derived from the error-prone bacteriophage promoters [11]. However, we have also used a linearized and purified RCA product to generate full-length infectious RNA transcripts from bacteriophage-driven clones, indicating the versatility of this system. Taken together, RCA represents a simple, high-fidelity, and cost-effective means to produce large amounts of plasmid DNA that can be repeatedly propagated and used to rescue infectious virus directly.

## 5. Acknowledgments

We would like to acknowledge VT-FAST, specifically Kristin Rose Jutras, Alexander Crookshanks, Michael Stamper, and Janet Webster, for there assistance in preparing this manuscript for publication.

We would thank members of the Weger-Lucarelli lab, specifically Tyler Bates, Emily Webb, and Pallavi Rai, for there valuable contributions and discussions in experimental design and manuscript preparation.

This work was partially supported by funding from DARPA’s PREventing EMerging Pathogenic Threats (PREEMPT) program.

